# NeuroMesh: A Bottleneck Topology Controller for Missing-Modality Brain Tumor Segmentation — A Mechanistic Pilot Study on BraTS

**DOI:** 10.64898/2026.08.23.746542

**Authors:** K Navaneetha Krishnan, K Janakiraman

## Abstract

Deep segmentation networks can degrade sharply when an expected MRI sequence is unavailable at inference. We present NeuroMesh, a bottleneck controller that combines a gated recurrent unit (GRU) with a graph-convolutional edge-activation mask, designed to adapt a U-Net-style segmentation backbone to missing input. We evaluate NeuroMesh in a pilot study using a 30-patient subset of the BraTS 2020 benchmark (22 training, 4 validation, and 4 held-out test patients) under a prespecified frozen-test protocol. On the frozen test set, NeuroMesh has higher tumor-core and enhancing-tumor Dice than a plain U-Net in most evaluated missing-modality conditions, but whole-tumor Dice falls from 0.596 to 0.108 when FLAIR is missing, compared with 0.604 to 0.545 for the plain U-Net. Direct analysis of the predicted edge-activation mask shows negligible change across modality-availability conditions. A parameter-light static-gating control reproduces the FLAIR failure mode without recurrence, a failure-signal input, or graph-structured machinery. These results do not support the intended interpretation that the trained controller performs input-conditional topology rewiring at the scale of this pilot. Instead, they expose a discrepancy between architectural intent and realized behavior and identify a specific missing-modality failure mode that warrants further investigation. Given the small validation and test sets, the findings are descriptive and do not establish clinical or population-level generalization.

## I. Introduction

**D**EEP convolutional networks achieve strong benchmark accuracy on multimodal brain tumor segmentation, but the accuracy figures reported in the literature are almost always obtained under an assumption that does not always hold at deployment: that all expected MRI sequences are present, co-registered, and of diagnostic quality. In the BraTS benchmark [16], [17], models are conventionally trained and evaluated using four structural sequences per patient — T1-weighted (T1), contrast-enhanced T1-weighted (T1ce), T2-weighted (T2), and T2 fluid-attenuated inversion recovery (FLAIR) — each sensitive to different tissue and pathological properties. T1ce is acquired after intravenous gadolinium-based contrast administration and is the only one of the four sequences that shows active blood–brain-barrier breakdown as a hyperintense enhancing rim; FLAIR suppresses the cerebrospinal fluid (CSF) signal and is comparatively sensitive to the vasogenic edema that surrounds a tumor. Because these four sequences are only partially redundant with one another radiologically, a network’s dependence on any single one is an empirical question, not something that can be assumed away. In practice, a sequence can be absent because it was not clinically indicated, failed acquisition or quality control, or was motion-corrupted, and a deployed model has to do something reasonable with the remaining channels rather than fail silently.

### A. Problem Statement

We consider missing-modality inference: one or more of the four input sequences is unavailable at inference time, and the network must still produce a usable segmentation from the remaining channels. This is distinct from, though related to, robustness to transient internal feature corruption (e.g. a hardware fault silencing a subset of feature-map channels at inference time), which the same controller design targets but which this pilot does not evaluate empirically; see Section VIII-D.

### B. Why Prior Work Is Insufficient

- **Dropout** [1] regularizes training but is fixed (or absent) at test time and does not adapt to which channels are *actually* missing for a given input; Monte Carlo dropout at test time is a common heuristic extension but still does not condition the dropout pattern on the true failure.
- **Ensembling** [2] can improve robustness but at proportional inference cost, without targeting the specific failure pattern present in a given input.
- **Mixture-of-experts and dynamic routing** [3], [4] adapt *computation* to input content but generally assume all routing targets remain available.
- **Modality-dropout training**, a common practical mitigation for missing-modality medical segmentation, trains a single network to tolerate channels missing during training but does not adapt its *internal connectivity* at inference time to the specific pattern of what is currently missing.

### C. Contribution

This paper makes the following contributions:

1. A bottleneck topology controller, NeuroMesh, combining a GRU-based recurrent state with a graph-convolutional edge-activation mask, designed to test whether a segmentation network can learn input-conditional internal rewiring in response to a missing-modality failure signal (Section III).
2. A pre-registered, single-touch evaluation protocol on a 30-patient BraTS 2020 subset, with a patient-level train/validation/test split frozen *before* any test-set evaluation, and every reported number traceable to a machine-readable result file (Section IV–V).
3. An empirical finding that NeuroMesh improves tumor-core (TC) and enhancing-tumor (ET) Dice over a plain U-Net baseline in this pilot, but exhibits a large, reproducible whole-tumor (WT) Dice collapse specifically when FLAIR is missing (Section VI).
4. A mechanistic analysis showing that the controller’s predicted edge-activation mask is not measurably conditioned on which modality is missing, that a parameter-light static-gating control reproduces the FLAIR failure mode, and that the “dynamic rewiring” mechanism motivating this architecture is therefore not empirically supported at the scale of this pilot (Section VII). We regard this negative mechanistic result, obtained through direct inspection rather than assumed, as the paper’s central contribution.

## II. Related Work

### A. Dynamic and Adaptive Neural Architectures

Mixture-of-experts and dynamic-routing methods [3], [4] adapt *computation* to input content but generally assume all routing targets (experts) remain healthy. NeuroMesh instead attempts to adapt *connectivity* in response to an explicit failure signal, a complementary (in principle) but empirically distinct goal from input-conditioned routing among always-available experts.

### B. Neural Architecture Search and Graph-Structured Networks

Differentiable architecture search [5] and graph neural architecture search [6] learn connectivity patterns at training time but fix them at deployment. Structured and lottery-ticket pruning [7] similarly yields a static sparse topology. NeuroMesh’s controller instead runs at inference time and is conditioned, at least in intent, on a per-input failure signal; Section VII examines whether this intent is realized in practice.

### C. Robustness in Medical Image Segmentation

Missing-modality robustness for multimodal MRI segmentation is a long-standing concern in the BraTS literature. Common approaches include modality-dropout training and modality-invariant feature learning; a direct, quantitative comparison against such missing-modality-specific baselines (e.g. HeMIS-style [14] hetero-modal architectures, or modality-invariant embedding methods) was not performed in this pilot and is identified as necessary future work (Section VIII-D) rather than implied by omission.

### D. Adversarial and Certified Robustness

Adversarial robustness [8] and certified defenses via randomized smoothing [9] address bounded *input* perturbations, not structural connectivity failures or missing input channels, and are not directly comparable baselines for the problem addressed here.

## III. Method

### A. Overview

NeuroMesh augments a 2D U-Net-style [11] encoder– decoder segmentation backbone with a topology controller inserted at the bottleneck. An overview of the complete architecture is shown in Fig. 1. The encoder maps a 4-channel input (T1, T1ce, T2, FLAIR; empirically verified channel order given in Section IV) through four downsampling stages to a bottleneck feature map *X*_5_ ∈ ℝ ^*B×C×h×w*^, where *C* = 16*b* for base channel width *b*. The bottleneck representation is globally pooled to obtain a feature summary, while a failure signal is computed from numerically inactive bottleneck channels. These signals are processed by the NeuroMesh controller, which comprises feature projections, a gated recurrent unit (GRU), and a graph controller that predicts an edge-activation mask. The mask modulates a learned base adjacency before the resulting bottleneck representation is further modulated by a sigmoid channel gate. The modified bottleneck is then passed to the U-Net decoder, which follows the standard upsampling-with-skip-connections pattern and produces per-pixel logits over *K* = 4 classes: background, necrotic/non-enhancing tumor core, peritumoral edema, and enhancing tumor, with BraTS label 4 remapped to contiguous index 3.

**Fig. 1.**
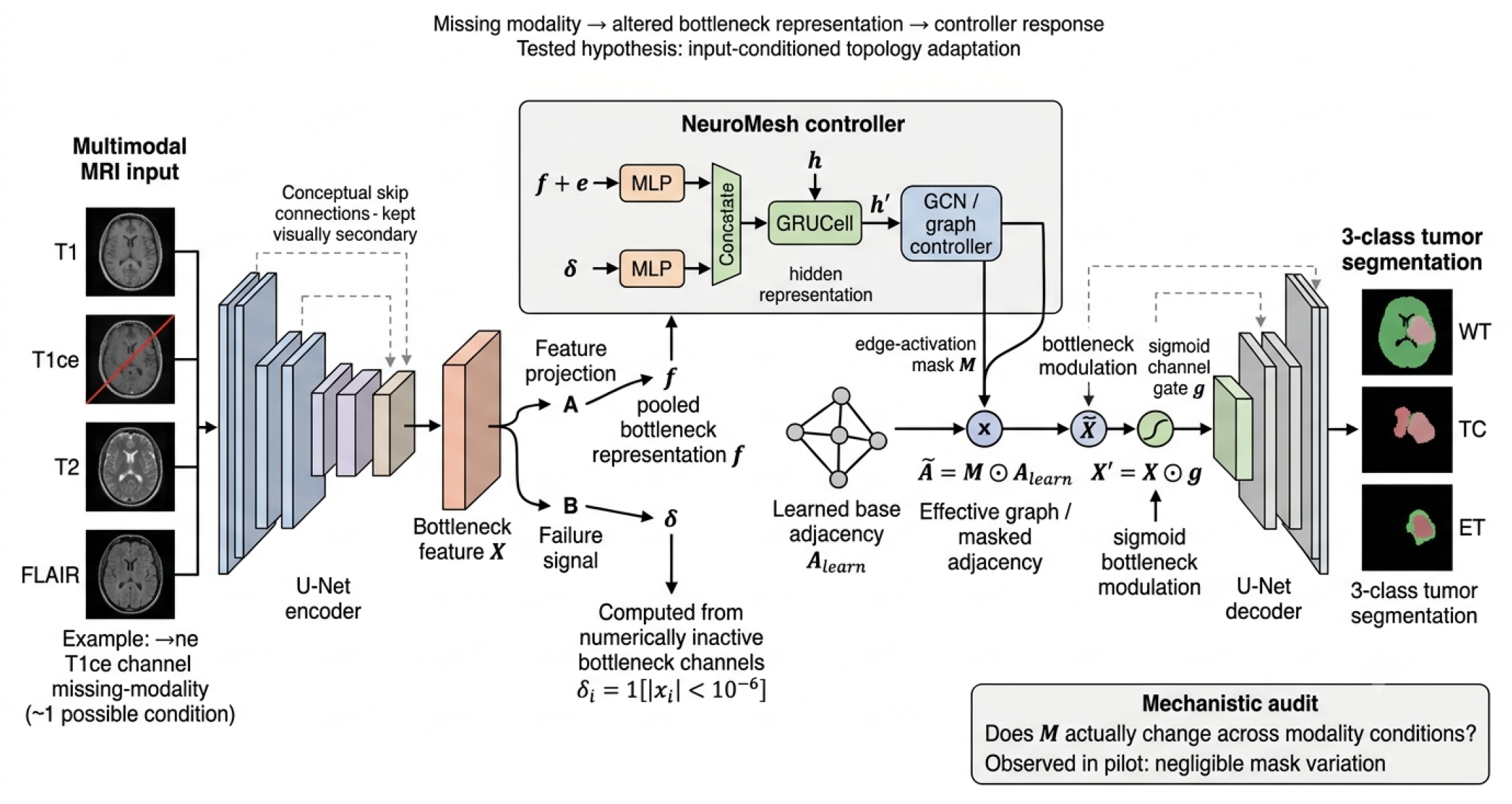
Overview of the NeuroMesh architecture. The 4-channel multimodal MRI input is processed by a U-Net encoder to obtain the bottleneck feature map ***X***_**5**_. A pooled bottleneck representation provides the feature input ***f***, while the failure signal ***δ*** is computed from numerically inactive bottleneck channels. The NeuroMesh controller combines these signals with the recurrent hidden state through MLP projections and a GRU cell, followed by a graph controller that predicts the edge-activation mask ***M*** . The effective adjacency is ***Ã* = *M* ⊙ *A***_**learn**_. The resulting bottleneck representation is subsequently modulated by a sigmoid channel gate ***g*** before being passed to the U-Net decoder. The figure illustrates the tested hypothesis of input-conditioned topology adaptation; the manuscript separately evaluates whether the predicted mask ***M*** actually changes across missing-modality conditions.

#### a) Ethical considerations

This study is a secondary analysis of publicly available, de-identified MRI data from the BraTS 2020 benchmark. No participants were recruited or contacted by the authors, and no new human-subject data were collected. The analysis used the benchmark data under its stated research-use terms.

### B. Graph Convolution Layer

Given node features 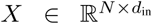 and a nonnegative adjacency *A* ∈ ℝ^*N×N*^ (self-loops included), the controller’s graph component uses the standard symmetric-normalized graph convolution of Kipf and Welling [12]:

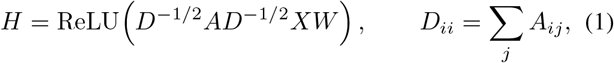

with 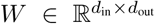 a learnable weight matrix and *D* the diagonal degree matrix (degrees clamped away from zero for numerical stability). In the controller, each of the *C* scalar entries of the pooled bottleneck vector is treated as a single graph node with a 1-dimensional feature (*d*_in_ = 1), which gives the layer a well-defined input without requiring an externally specified graph structure.

### C. Topology Controller

Let *x* ∈ ℝ^*C*^ denote the global-average-pooled bottleneck feature vector for a given input (batch dimension omitted for clarity), 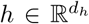 a recurrent hidden state carried across forward passes, and *δ* ∈ { 0,1}^*C*^ a binary failure signal. Unless supplied externally, *δ* is derived automatically as

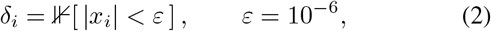

i.e. a channel is flagged as failed if it is numerically dead, which is intended as an automatic, architecture-internal proxy for a missing or corrupted upstream signal. The controller then computes

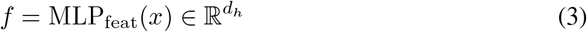

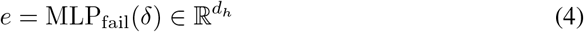

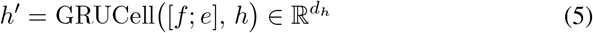

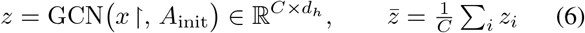

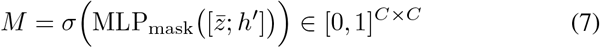

where *x* ∈ ∈ ℝ^*C×*1^ denotes *x* reshaped into *C* scalar-feature nodes, [· ; ·] denotes concatenation, *σ* is the logistic sigmoid, and *A*_init_ ∈ ℝ^*C×C*^ is a learnable adjacency matrix initialized to the identity, so that the controller begins training close to the identity transform (no rewiring) and learns off-diagonal connectivity from there. During training, *M* is produced with a straight-through Bernoulli estimator [13] — a hard sample 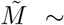 Bernoulli(*M*) is used on the forward pass, while gradients are backpropagated through the soft sigmoid, *M* ← *M* + sg 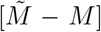, with sg[·] the stop-gradient operator; at inference, *M* is used directly as the deterministic clipped sigmoid. The controller output is

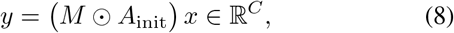

i.e. *y*_*i*_ = ∑_*j*_ *M*_*ij*_ *A*_init,*ij*_ *x*_*j*_, a masked linear combination of the pooled bottleneck channels under the (learnable, maskgated) adjacency.

#### Algorithm 1

NeuroMesh controller forward pass (single step)

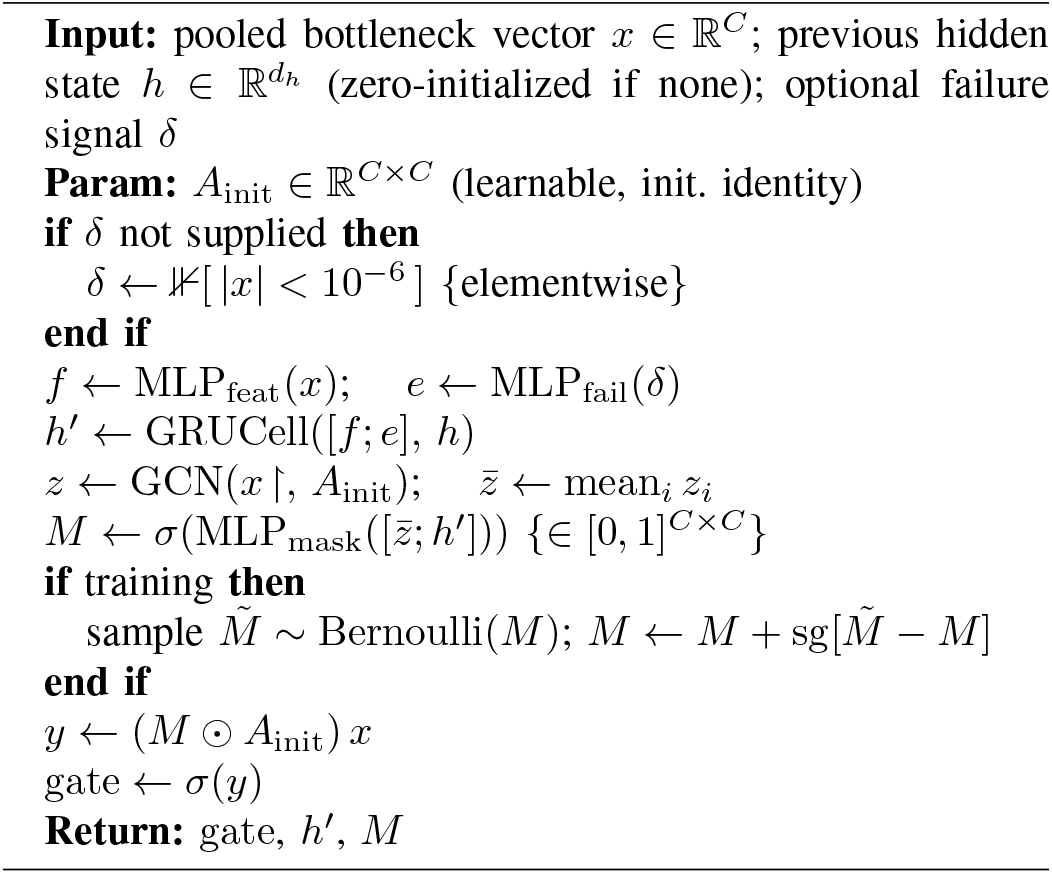

### D. Segmentation Backbone

The controller’s output *y* ∈ ℝ^*C*^ is passed through a sigmoid to form a per-channel gate, gate = *σ*(*y*) ∈ [0, 1]^*C*^, which is broadcast spatially and applied multiplicatively to the bottle-neck feature map, 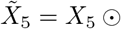 gate, before decoding. At *b* = 16 (the configuration used in this pilot’s reported experiments; Section V), *C* = 256, so the mask *M* has *C*^2^ = 65,536 entries, which dominates the controller’s parameter count relative to a plain channel-attention gate of the same width.

### E. Composite Training Objective

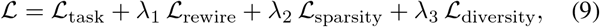

with (*λ*_1_, *λ*_2_, *λ*_3_) = (0.1, 0.01, 0.001) throughout this pilot —these are the architecture’s originally proposed defaults, used unmodified, and are not claimed to be tuned or optimal (Section VIII-D).

#### Task loss

An equal-weighted combination of pixelwise cross-entropy and soft multi-class Dice loss over the *K* = 4 classes:

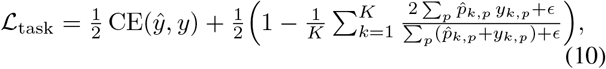

where 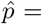 softmax 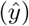, *p* indexes pixels, *y*_*k,p*_ is the one-hot ground truth, and *ϵ* = 10^*−*6^.

#### Rewiring loss

A temperature-scaled (*T* = 2) Kullback– Leibler divergence between spatially pooled logits on the clean input and on a modality-corrupted version of the same input, encouraging the controller to find a rewiring under which the predictive distribution is preserved despite the corruption:

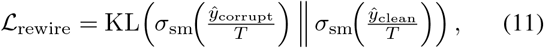

with *σ*_sm_ the softmax and logits spatially averaged before the divergence is computed.

#### Sparsity loss

The fraction of active (> 0.5) entries of the predicted mask, averaged over *L* forward passes in a training batch, encouraging a sparse rewiring:

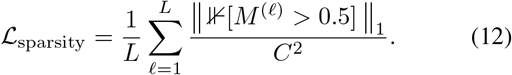

#### Diversity loss

A hinge penalty (margin 0.5) on pairwise cosine similarity between flattened masks from different forward passes in the same batch, discouraging a degenerate, input-independent policy:

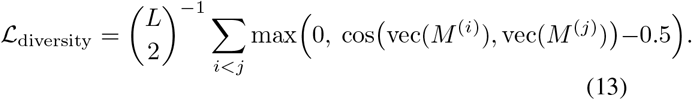

This term is, in effect, an explicit training-time penalty for the very failure mode Section VII documents empirically; that it did not prevent the near-degenerate solution observed is itself informative and is discussed in Section VIII.

### F. On a Closed-Form Robustness Bound

An earlier stage of this project considered stating a closed-form bound on expected output error under random node failures, of the schematic form E[∥*z*_rewired_ −*z*_clean_∥ ^2^] *≲* (*Kp* + *p/*(1 − (1− *ρ*)^2^) ) ∥*x*∥^2^ for Lipschitz constant *K*, node-failure rate *p*, and mask density *ρ*. **No such bound is proven or asserted in this paper**. The supporting argument available at that stage relied on an unstated concentration-inequality step and did not establish that the learned mask *M* achieves the assumed redundancy in practice; given the mechanistic finding of Section VII — that *M* is close to input-invariant in this pilot’s trained model — any such bound would in any case describe a regime the trained model does not appear to occupy. A rigorous derivation, with explicit probabilistic assumptions, is left to future work (Section VIII-D) rather than presented here in unproven form.

## IV. Data

### A. Dataset and Provenance

This pilot uses a random 30-patient subset (seed = 42) of the BraTS 2020 training set [16]–[18], obtained via the pre-sliced HDF5 mirror distributed on Kaggle as `awsaf49/brats2020-training-data` rather than the official CBICA NIfTI release. Each of the 369 full-set volumes contains 14.8–65.8% tumor-visible slices, so an unconditioned random draw is not expected to introduce a tumor-size selection bias, though this was not independently verified against the specific 30-patient draw beyond that population-level bound. Each patient volume comprises four co-registered, skull-stripped MRI sequences of native resolution 240 × 240 over 155 axial slices, with an expert segmentation label volume using labels { 0, 1, 2, 4 } (background, necrotic/non-enhancing tumor core, peritumoral edema, enhancing tumor), with label 4 remapped to a contiguous index 3 for training, consistent with standard BraTS practice.

### B. Modality Channel-Order Verification

The HDF5 mirror does not document which of its four image channels corresponds to which MRI sequence. This was determined empirically, not assumed, via three independent checks on real downloaded slices: (1) a region-contrast test, comparing mean intensity inside each tumor sub-region against surrounding tissue, which showed a large, region-specific enhancement jump on one channel matching the expected gadolinium-contrast signature of T1ce; (2) direct visual inspection of CSF/ventricle appearance, which is bright (hyperintense) on FLAIR and dark (suppressed) on the other three sequences among these four; and (3) inspection of the choroid plexus, a structure that normally shows contrast enhancement on post-gadolinium imaging, which was visible specifically on the channel identified as T1ce by check (1). The three checks converge on

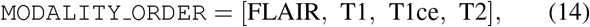

which we adopt with reasonable but not certain confidence: this is the authors’ own visual/statistical determination on a sample of slices, not documentation from the dataset publisher, and we flag it as such rather than stating it as an established fact.

### C. Patient-Level Splitting

To avoid evaluation leakage from the 155 highly correlated adjacent slices belonging to one patient appearing across splits, the 30-patient subset was partitioned at the *volume* level, before any slice-level indexing, using a deterministic seeded shuffle (`val_fraction`= 0.15, `test_fraction`= 0.15, `seed`= 42), giving:

- **Train** (*n* = 22): patient IDs 13, 14, 16, 17, 45, 58, 72, 102, 113, 120, 141, 215, 217, 230, 280, 288, 303, 309, 328, 347, 360, 367.
- **Validation** (*n* = 4): patient IDs 112, 126, 259, 333.
- **Test** (*n* = 4, frozen): patient IDs 48, 53, 115, 302.

The test split was frozen *before* any evaluation of it, under a written protocol (Section V-C). One minor, disclosed deviation from strict isolation occurred: the modality-order verification above used slices from patients 45, 48, and 141 (raw image appearance only, no labels), and patient 48 is a test-split patient. No label information or model-relevant signal was used from patient 48 in that check — modality identity is a fixed scanner/protocol property, not a property of an individual patient’s pathology — but strictly, zero test-patient pixels should have been examined before the freeze, and this is recorded rather than omitted.

### D. Preprocessing

Each 2D slice is normalized per channel to zero mean and unit variance. Native 240× 240 images are bilinearly downsampled, and label maps nearest-neighbor downsampled, to 96 ×96 prior to training and evaluation; predictions are nearest-neighbor upsampled back to 240 ×240 before metrics are computed against the native-resolution ground truth. This resolution reduction is a compute-budget accommodation for the single-CPU-core hardware this pilot ran on (Section V), not a methodological choice, and is treated as a limitation (Section VIII-D) rather than a design decision to be re-derived at native resolution before any number here is treated as final.

### E. Modality-Dropout Protocol

At training time, each input channel is independently zeroed with probability *p* = 0.15. At evaluation time, either a specific channel is deterministically zeroed for all patients in a given condition (single-modality-missing conditions, used for all results in this paper) or left unmodified (clean condition); randomized multi-channel dropout sweeps and correlated/cascading failure models were scoped out of this pilot’s protocol (Section V-C) and are not reported.

## V. Experimental Setup

### A. Models Compared

- **NeuroMesh** (proposed): the full architecture of Section III, base channel width *b* = 16 (19,149,108 total parameters).
- **Plain U-Net**: identical backbone, no bottleneck controller.
- **DropoutUNet**: identical backbone with Monte Carlo dropout at the bottleneck.
- **StaticGatedUNet**: identical bottleneck *gating operation* as NeuroMesh, but the gate is produced by a plain feedforward MLP on the pooled bottleneck vector, with no recurrence, no failure-signal input, and no learned adjacency mask (≈ 1.98 ×10^6^ parameters, i.e. only on the order of 3 × 10^4^ more than Plain U-Net). This model isolates the contribution of “a learned bottleneck gate exists at all” from the recurrent/graph controller machinery specific to NeuroMesh.
- **EnsembleUNet**: implemented and unit-tested but *not* trained or evaluated on real data in this pilot; excluded from all results below.

### B. Training Configuration

All four trained models used AdamW [15] (learning rate 10^*−*3^, weight decay 10^*−*5^), batch size 8, for 4 epochs (1,708 optimizer steps: 427 batches/epoch × 4) over the 22-patient training split, with `torch.manual_seed(0)`. Training-time modality dropout used *p* = 0.15 per channel, independent across channels. NeuroMesh’s composite loss used the default weights of Eq. (9). Training ran on **1 CPU core, 4 GB RAM, no GPU** (PyTorch 2.13 CPU build, Python 3.12, SciPy 1.17, Ubuntu container); this hardware constraint, disclosed rather than a design choice, motivated both the 96 × 96 training resolution (Section IV) and a checkpoint/resume training loop (each invocation trained until a hard *∼* 280–380 s wall-clock limit and saved state; the reported NeuroMesh checkpoint required 13 resumed invocations, 1,950 s cumulative wall-clock). DataLoader shuffling order was not separately seeded per resume chunk, so exact batch order is not bitwise-reproducible from a from-scratch rerun, though results are expected to be statistically similar given the same weight-initialization seed. A single additional checkpoint, *neuromesh e8*, continued training the frozen NeuroMesh checkpoint for 4 further epochs (8 total) with optimizer momentum/variance state reset at the 4-epoch boundary (the original checkpoint did not persist optimizer state); *neuromesh e8* was evaluated on validation only and is used solely for the epoch-4-vs-epoch-8 mechanistic comparison in Section VII.

### C. Frozen Evaluation Protocol

Before any test-set evaluation, a written protocol (`PROTOCOL_FREEZE_v1.md`) fixed the data split, model configuration, checkpoint (identified by SHA-256 checksum), and the exact set of metrics and conditions to be evaluated on test, with the explicit rule that test-set results would be recorded as obtained, with no retraining, hyperparameter change, or architecture change permitted in response to them under this protocol version; any such change would require a new, independently frozen protocol version evaluated against its own, still-untouched test split. Under this protocol, only NeuroMesh and Plain U-Net were evaluated on the frozen test set; DropoutUNet and StaticGatedUNet results reported below are validation-set only, and are explicitly labeled as such in every table.

### D. Metrics

For each patient and each tumor sub-region — whole tumor (WT = {1, 2, 3}), tumor core (TC = {1, 3}), enhancing tumor (ET = {3}), following standard BraTS grouping — we compute the Dice similarity coefficient,

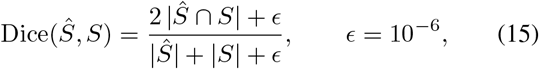

the 95th-percentile symmetric Hausdorff distance (HD95, via boundary erosion and a Euclidean distance transform, assuming 1 mm isotropic spacing per BraTS convention — not independently re-verified against this specific mirror, which does not carry NIfTI affine metadata), and region-wise sensitivity and precision, each computed on the *full reconstructed volume* (155 native-resolution slices stacked in *z*-order, not slice-averaged), matching how BraTS itself scores challenge submissions. HD95 and precision are reported as undefined (not zero, not infinite) when the corresponding region is empty in ground truth or prediction. Given *n* = 4 per split, we report descriptive statistics (mean ± SD across patients) throughout and explicitly do not compute inferential statistics; see Section VI-E.

## VI. Results

### A. Frozen Test-Set Results: NeuroMesh vs. Plain U-Net

Table I reports Dice (mean ± SD, *n* = 4 test patients: 48, 53, 115, 302) and mean HD95 for both models evaluated under this protocol. NeuroMesh exceeds Plain U-Net on TC in three of five conditions and on ET in four of five conditions, but its WT Dice under FLAIR-missing (0.108, SD 0.100) is markedly worse than Plain U-Net’s under the same condition (0.545, SD 0.240) and worse than NeuroMesh’s own clean-input WT Dice (0.596), a Δ = −0.488 absolute drop versus Plain U-Net’s Δ = −0.059 under the same channel loss.

**TABLE I.** Frozen test-set results (***n* = 4** patients: 48, 53, 115, 302), NeuroMesh vs. Plain U-Net. Dice reported as mean **±** SD across patients; HD95 reported as the mean across patients for which both masks were non-empty (see Section v); “n/a” indicates the region was empty in every test patient’s prediction under that condition. Source: `results/frozen`/FROZEN_TEST_`{neuromesh,plain}_real30.json`.

| Condition | Model | Dice |  |  | HD95 (mm) |  |  |
| --- | --- | --- | --- | --- | --- | --- | --- |
|  |  | WT | TC | ET | WT | TC | ET |
| Clean (all 4) | NeuroMesh | 0.596 ± 0.259 | 0.462 ± 0.255 | 0.560 ± 0.117 | 15.7 | 43.5 | 52.6 |
| | Plain U-Net | 0.604 ± 0.235 | 0.230 ± 0.229 | 0.396 ± 0.232 | 23.0 | 61.2 | 51.8 |
| FLAIR missing | NeuroMesh | **0.108 ± 0.100** | 0.219 ± 0.215 | 0.357 ± 0.210 | 38.5 | 63.8 | 72.7 |
| | Plain U-Net | 0.545 ± 0.240 | 0.177 ± 0.207 | 0.364 ± 0.250 | 60.3 | 78.4 | 69.1 |
| T1 missing | NeuroMesh | 0.595 ± 0.259 | 0.389 ± 0.194 | 0.410 ± 0.104 | 21.4 | 52.8 | 51.2 |
| | Plain U-Net | 0.509 ± 0.256 | 0.173 ± 0.215 | 0.209 ± 0.197 | 21.5 | 41.1 | 46.5 |
| T1ce missing | NeuroMesh | 0.504 ± 0.359 | 0.003 ± 0.005 | $\approx 0$ | 15.4 | 26.5 | n/a |
| | Plain U-Net | 0.512 ± 0.321 | $\approx 0$ | $\approx 0$ | 13.8 | n/a | n/a |
| T2 missing | NeuroMesh | 0.509 ± 0.327 | 0.447 ± 0.264 | 0.570 ± 0.123 | 18.7 | 55.5 | 63.6 |
| | Plain U-Net | 0.601 ± 0.212 | 0.215 ± 0.233 | 0.380 ± 0.234 | 26.8 | 68.3 | 59.4 |

### B. Development-Time Four-Model Comparison

Table II extends the comparison to DropoutUNet and StaticGatedUNet, which were evaluated on the validation split only (patients 112, 126, 259, 333) and are not part of the frozen protocol; they are reported for mechanistic context (Section VII), not as claims of superiority or inferiority against the frozen numbers above, since validation and test are different patients. On validation, the FLAIR-missing WT collapse pattern is reproduced by both NeuroMesh (0.255, SD 0.260) and StaticGatedUNet (0.228, SD 0.234), while Plain U-Net (0.666) and DropoutUNet (0.664) do not show it.

**TABLE II.** Development-time validation comparison across four models (***n* = 4** patients: 112, 126, 259, 333). Dice reported as mean **±** SD. **V****alidation split, not the frozen test set**; DropoutUNet and StaticGatedUNet were not evaluated on test under this protocol version (Section V-C). Source:`results/manifests/four_modelvalidation.md`, regenerated directly from `results/validation`/VAL_*_`regions.json`.

| Condition | Model | WT Dice | TC Dice | ET Dice |
| --- | --- | --- | --- | --- |
| Clean | NeuroMesh (4 ep.) | 0.718 ± 0.224 | 0.360 ± 0.343 | 0.396 ± 0.319 |
| | Plain U-Net | 0.788 ± 0.057 | 0.252 ± 0.246 | 0.299 ± 0.283 |
| | DropoutUNet | 0.833 ± 0.059 | 0.458 ± 0.251 | 0.370 ± 0.346 |
| | StaticGatedUNet | 0.691 ± 0.272 | 0.348 ± 0.334 | 0.394 ± 0.356 |
| FLAIR missing | NeuroMesh (4 ep.) | **0.255 ± 0.260** | 0.367 ± 0.367 | 0.374 ± 0.375 |
| | Plain U-Net | 0.666 ± 0.085 | 0.288 ± 0.289 | 0.329 ± 0.325 |
| | DropoutUNet | 0.664 ± 0.065 | 0.481 ± 0.172 | 0.367 ± 0.359 |
| | StaticGatedUNet | **0.228 ± 0.234** | 0.346 ± 0.346 | 0.361 ± 0.361 |
| T1 missing | NeuroMesh (4 ep.) | 0.817 ± 0.054 | 0.419 ± 0.221 | 0.425 ± 0.236 |
| | Plain U-Net | 0.658 ± 0.120 | 0.277 ± 0.158 | 0.305 ± 0.172 |
| | DropoutUNet | 0.798 ± 0.053 | 0.500 ± 0.233 | 0.438 ± 0.266 |
| | StaticGatedUNet | 0.698 ± 0.153 | 0.369 ± 0.251 | 0.383 ± 0.270 |
| T1ce missing | NeuroMesh (4 ep.) | 0.616 ± 0.348 | 0.0002 ± 0.0004 | $\approx 0$ |
| | Plain U-Net | 0.738 ± 0.112 | $\approx 0$ | $\approx 0$ |
| | DropoutUNet | 0.820 ± 0.065 | 0.447 ± 0.276 | $\approx 0$ |
| | StaticGatedUNet | 0.659 ± 0.299 | 0.378 ± 0.218 | 0.0005 ± 0.0009 |
| T2 missing | NeuroMesh (4 ep.) | 0.625 ± 0.325 | 0.359 ± 0.353 | 0.395 ± 0.324 |
| | Plain U-Net | 0.751 ± 0.082 | 0.239 ± 0.229 | 0.294 ± 0.257 |
| | DropoutUNet | 0.829 ± 0.063 | 0.471 ± 0.266 | 0.387 ± 0.357 |
| | StaticGatedUNet | 0.764 ± 0.138 | 0.337 ± 0.325 | 0.389 ± 0.353 |

### C. Patient-Level Breakdown of the WT/FLAIR Failure

Fig. 2 shows per-patient WT Dice under FLAIR-missing on validation. The failure is not uniform across the four patients: NeuroMesh and StaticGatedUNet collapse to near-zero Dice (< 0.01) specifically on patients 259 and 333, while retaining Dice comparable to (patient 112) or exceeding (patient 126) the other two models on the remaining two patients. This patient-specific, bimodal pattern — rather than a uniform, graded degradation — is consistent with a small number of cases where the model’s WT prediction collapses toward an empty (or near-empty) mask once FLAIR is removed, rather than a smoothly reduced-quality segmentation.

**Fig. 2.**
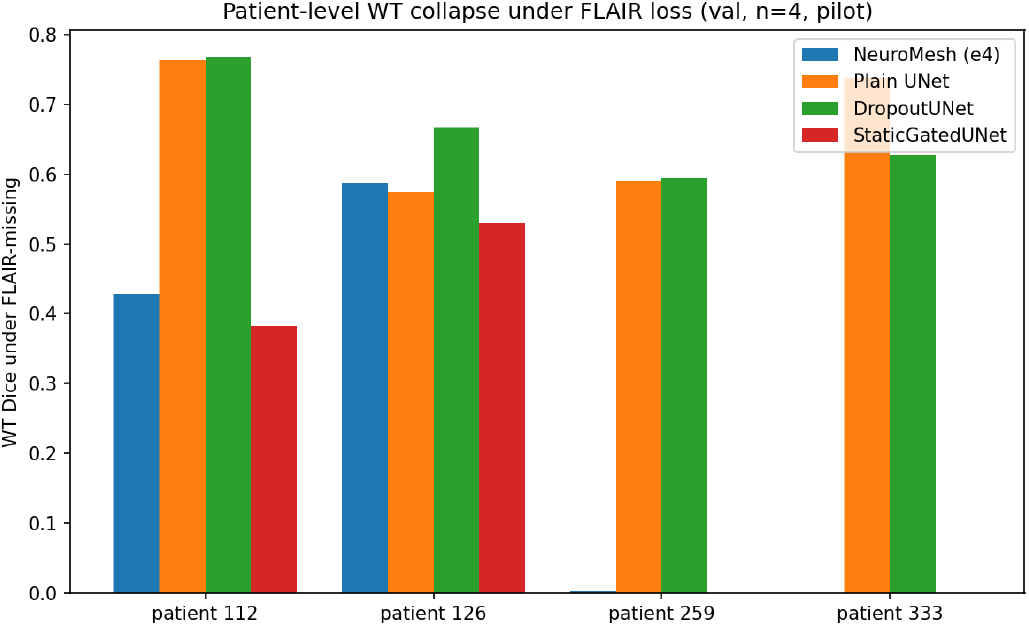
per-patient whole-tumor (wt) dice under flair-missing, validation split (***n* = 4**). neuromesh and staticgatedunet collapse to near-zero dice on patients 259 and 333 specifically, while plain u-net and dropoutunet do not. source: `results/manifests/patient_level_flair missing_wt.md.`

### D. The Enhancing-Tumor/T1ce Ceiling

Under T1ce-missing, ET Dice is ≈ 0 for *every* model evaluated, including both baselines (Tables I–II). Because ET is defined by the presence of gadolinium contrast enhancement, and T1ce is the only one of the four sequences that carries this contrast signal, this pattern is consistent with an information-availability ceiling common to all architectures tested, rather than an architecture-specific weakness of any one model; TC Dice under the same condition similarly collapses toward zero for NeuroMesh and Plain U-Net, though less completely for DropoutUNet and StaticGatedUNet (Table II), a difference we do not have a mechanistic explanation for and are not offering a post-hoc one.

### E. Statistical Considerations

With *n* = 4 test patients and *n* = 4 validation patients, this pilot does not support inferential statistical testing (e.g. a paired *t*-test or Wilcoxon signed-rank test across patients), and none is reported; differences above are described descriptively. For context: a two-sample *t*-test detecting a moderate-to-large effect (Cohen’s *d* ≈ 0.8) at two-sided *α* = 0.05 and 80% power requires, by the standard normal-approximation sample-size formula *n* ≈ 2(*z*_1*−α/*2_ + *z*_1*−β*_)^2^*/d*^2^, on the order of 25 patients per arm — roughly six times this pilot’s test-set size. The WT/FLAIR effect reported above (Δ ≈ −0.49 Dice for NeuroMesh vs. Δ ≈ −0.06 for Plain U-Net) is large relative to the observed patient-to-patient SD and reproduces with the same direction and a similar magnitude on both the validation and the frozen test split (independent patient sets), which is more reassuring than a single-split result of this size would be in isolation, but *n* = 4/*n* = 4 remains a pilot scope, not a basis for a generalization claim.

## VII. Mechanistic Analysis of the Controller

The results above establish that NeuroMesh differs from a plain U-Net in this pilot. This section asks *why*, by directly instrumenting the trained controller rather than assuming its intended mechanism (input-conditional topology rewiring) is what produced the observed differences.

### A. The Mask Is Not Measurably Input-Dependent

For every (patient, condition) pair checked, at both 4 and 8 epochs of training, the mean absolute difference between the predicted mask *M* under clean input and under any single-modality-missing condition was ≈0.00000, and the fraction of the *C*^2^ = 65,536 mask entries changing by more than 0.05 between clean and any failure condition was 0.0000 for every patient checked (Table III). This held at both checkpoints and, if anything, the already-negligible difference shrank further between epoch 4 and epoch 8 rather than growing with additional training. Visual inspection of rendered mask heatmaps (available in the accompanying repository) shows an unstructured pattern that is visually indistinguishable across all five conditions. This directly contradicts input-conditional rewiring as an established property of this trained model: whatever the mask MLP has learned, it is not conditioning meaningfully on which modality is present or absent.

**Table III.**
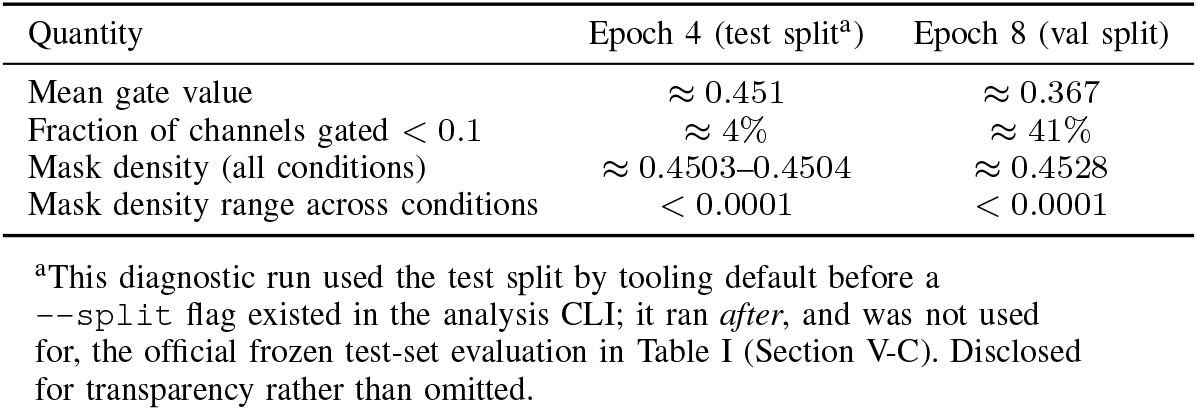
controller mechanism summary, mean across the 4 patients checked at each epoch. “Mask density” is the mean fraction of active (**> 0.5**) mask entries. Gate statistics are for the sigmoid-activated channel gate applied to the bottleneck. Source:`results/mechanism/epoch{4,8}_*/mechanism_summary.json`.

### B. Gate Selectivity Increased With Training, but Not Conditionally

Between epoch 4 and epoch 8 (same 22 training patients, continued training, no architecture change), the mean gate value dropped from ≈ 0.451 to ≈ 0.367, and the fraction of the *C* = 256 bottleneck channels strongly suppressed (gate < 0.1) rose from ≈ 4% to ≈ 41% (Table III). This shift is nearly identical whether the input is clean or has a modality missing: the gate became a substantially more selective *fixed* filter with additional training, not a more *conditional* one. One partial exception is notable: under T1ce-missing specifically, the fraction of strongly suppressed channels at epoch 4 was lower (≈ 0.7%) than under any other condition, consistently across all four patients checked — a real, reproducible, condition-specific signal, but a small one in absolute terms, and one that corresponds to the controller becoming *less* selective, not more, exactly where TC/ET performance collapses toward zero under that same condition (Section VI). We report this pattern without a mechanistic explanation for it.

### C. A Parameter-Light Static Control Reproduces the Failure Mode

StaticGatedUNet has the same bottleneck gating *operation* as NeuroMesh but no recurrence, no failure-signal input, and no learned adjacency mask; it is not parameter-matched to the full controller (≈ 1.98 × 10^6^ parameters vs. NeuroMesh’s (19.1) 10×6, a gap dominated by the mask MLP’s *C*× *C* output layer specifically excluded by this control’s design).

Despite this, StaticGatedUNet collapses on the *same two validation patients*, under the *same condition* (FLAIR-missing, WT specifically), as NeuroMesh does, almost exactly (Fig. 2). This is consistent with NeuroMesh’s WT/FLAIR fragility arising from the presence of a learned bottleneck gate as such, rather than from anything specific to the controller’s recurrent or graph-structured machinery. It does *not*, however, reproduce NeuroMesh’s TC recovery under T1ce-missing (StaticGate: 0.378; NeuroMesh: ≈0; Plain: ≈ 0; DropoutUNet: 0.447; Table II) — this particular pattern groups {DropoutUNet, StaticGatedUNet} against {Plain U-Net, NeuroMesh}, which does not split cleanly along “has a bottleneck gate,” and we do not offer a post-hoc explanation for it.

### D. Synthesis: What Can and Cannot Be Claimed

#### Can claim, at the scale of this pilot

NeuroMesh’s TC/ET advantage over Plain U-Net is real and fairly consistent across conditions; its WT/FLAIR fragility is real, reproducible across independent patient sets (validation and frozen test), and closely mirrored by a much simpler static-gating control; the controller’s predicted mask has not developed measurable input-dependent structure at the training scale used here (22 training patients, up to 8 epochs, 96 × 96 resolution). **Cannot claim:** that NeuroMesh performs dynamic, input-conditional topology adaptation; that its performance differences from Plain U-Net arise specifically from the recurrent or graph-structured controller machinery rather than from the mere presence of a learned bottleneck gate; that any of the above generalizes beyond the 4 validation and 4 test patients evaluated; or that substantially longer training would, or would not, eventually produce condition-dependent mask structure — only 4 and 8 epochs were checked.

## VIII. Discussion

### A. Biological Interpretation of the Modality-Specific Failure Modes

The two dominant empirical patterns in this pilot both have a plausible grounding in MRI physics rather than requiring an architecture-specific explanation. FLAIR suppresses CSF signal and is comparatively the most sensitive of the four sequences to peritumoral vasogenic edema, which is the largest single component (by volume, in most gliomas) of the WT label group; removing FLAIR is therefore expected *a priori* to disproportionately harm WT delineation relative to the other three sequences, and does so for every model tested here to some degree (Table I), with NeuroMesh and StaticGatedUNet showing a substantially larger effect than Plain U-Net and DropoutUNet. Conversely, ET is defined operationally by gadolinium contrast enhancement, visible only on T1ce among these four sequences; its collapse to ≈0 Dice under T1ce-missing for all four models tested is consistent with a genuine information ceiling rather than a fixable modeling deficiency, and should not, on this evidence, be read as an architecture comparison at all.

### B. Intended vs. Observed Operating Mechanism

NeuroMesh was designed to test whether a segmentation network could learn to *rewire itself*, in an input-conditional way, in response to a missing or corrupted imaging modality. The mechanistic analysis of Section VII does not support that this occurred in the trained model evaluated here: the predicted mask is close to input-invariant, and a simpler static control reproduces much of the same behavior. A largely static, learned bottleneck transformation is, on the evidence available, a more defensible description of what this pilot’s trained NeuroMesh model actually computes than “dynamic topology rewiring.” We regard reporting this distinction plainly, rather than describing NeuroMesh only in terms of its intended mechanism, as more valuable to the field than the raw Dice comparisons in Section VI on their own.

### C. Open Questions

This pilot raises, without resolving, three questions for follow-up work: (1) whether mask differentiation would emerge with substantially more training, or whether a fully unconstrained *C* × *C* = 65,536-parameter mask is simply too large a search space for 22 training patients to constrain regardless of epoch count; (2) what explains the TC/T1ce-missing pattern that groups {DropoutUNet, StaticGatedUNet} against {Plain U-Net, NeuroMesh}, which cuts against the simplest “it’s just the gate” account of Section VII; and (3) whether a smaller, more constrained mask (e.g. a low-rank or block-structured adjacency) would learn condition-dependent structure more readily than the current fully free formulation.

### D. Limitations

1. **Pilot sample size**. *n* = 4 validation, *n* = 4 test patients. This is a pilot, not a generalization study (Section VI-E); no claim in this paper should be read as established for the broader BraTS population or for any clinical population.
2. **Reduced training/evaluation resolution**. 96 × 96, against a native BraTS resolution of 240 ×240, a compute accommodation (Section IV) rather than a methodological choice. Re-running at native resolution on adequate compute is the natural first step before treating any number in this paper as final.
3. **Single random seed**. All four trained models used one training seed; no seed-variance estimate is available.
4. **Unvalidated loss weights**. (*λ*_1_, *λ*_2_, *λ*_3_) in Eq. (9) are carried-over defaults, not validated against held-out Dice (Section III-E).
5. **No formal robustness bound**. See Section III, final subsection; only an empirical robustness curve (Section VI) is reported.
6. **Corrupted-feature-channel robustness not evaluated**. The controller’s failure signal (Eq. (2)) is designed to generalize to transient internal feature corruption, not only missing input modalities, but this pilot’s evaluation protocol (Section V-C) covers only missing-modality conditions; correlated, cascading, and Byzantine failure-model evaluation is implemented in the accompanying codebase but was explicitly scoped out of this protocol version and not run.
7. **EnsembleUNet not evaluated**. Implemented and unit-tested only.
8. **Modality-order provenance**. The channel-order mapping (Section IV) is the authors’ own empirical determination, not documentation from the dataset publisher, albeit corroborated by three converging independent checks.
9. **Missing-modality-specific baselines not compared**. No comparison against hetero-modal architectures (e.g. HeMIS-style [14]) or modality-invariant feature-learning methods specific to the missing-modality BraTS literature (Section II).

### E. Ethical and Data-Use Considerations

This work uses the publicly available, de-identified BraTS dataset under its associated citation and data-use terms [16]– [18]; no additional institutional patient data was used or referenced at any point in this project. As BraTS is a publicly released, de-identified research dataset, no institution-specific IRB/ethics approval applies to its use in this form; readers intending to apply this or a derived model to any non-public or identifiable clinical data should obtain the appropriate institutional and regulatory approvals independently. NeuroMesh, as evaluated here, is a research prototype and is not validated for, and must not be used for, clinical diagnosis, treatment planning, or patient care.

## IX. Reproducibility Statement

All primary experimental results reported in this paper are traceable to machine-readable JSON or CSV files in the accompanying code release. These files record the command-line invocation, dataset split sizes, patient identifiers, compute device, wall-clock time, timestamp, and, for the frozen test-set checkpoints, a SHA-256 checksum. A provenance manifest (`results/manifests/RESULTS_MANIFEST.md`) documents the source of every numerical result reported in Tables I–III and Fig. 2.

The frozen NeuroMesh checkpoint is identified by the following SHA-256 checksum: `19108deeffbca2723a4edf9dc046164712b5b69decbe98d7bf88 4448af152962`

The train/validation/test split (Section IV-C) is fully specified by patient identifier and is reproducible from the public metadata CSV using the stated seed. Software dependencies are pinned in `pyproject.toml`. The automated test suite uses synthetic fixtures only and requires no licensed data; it is executed through continuous integration and verifies tensor shapes, gradient flow, loss finiteness, metric correctness, and a regression test for a hidden-state/batch-size mismatch identified and corrected during real-data evaluation.

These automated tests verify software behavior but do not verify the real-data scientific results in Section VI, which require a manual rerun using the licensed dataset via `neuromesh reproduce`. Code, configuration, and frozen results are provided through the repository and archived release identified in the Data Availability statement. The archived release should be used as the reference version for reproducing the specific results reported in this manuscript. No manuscript DOI had been assigned at the time of writing.

## X. Conclusion

We presented NeuroMesh, a GRU- and graph-convolution-based bottleneck controller for missing-modality-robust brain tumor segmentation, together with a fully instrumented, pre-registered pilot evaluation on a 30-patient BraTS 2020 subset. In this pilot, NeuroMesh improves tumor-core and enhancing-tumor Dice over a plain U-Net baseline under most missing-modality conditions, but exhibits a large, reproducible whole-tumor Dice collapse when FLAIR is missing. Direct mechanistic inspection shows that this behavior does not arise from the input-conditional topology rewiring the architecture was designed to achieve: the controller’s predicted mask is close to input-invariant, and a parameter-light static-gating control reproduces the same failure mode. We therefore present NeuroMesh’s empirical performance profile and its mechanistic non-explanation together, as we consider the latter the more durable contribution of this pilot. Future work should prioritize, in order: re-running this protocol at native resolution on adequate compute; scaling patient count sufficiently to support inferential statistics (Section VI-E indicates on the order of 25 patients per arm for a moderate effect); completing the EnsembleUNet and correlated/cascading failure-model evaluations already implemented but unrun; comparing against missing-modality-specific baselines from the BraTS literature (Section II); and, if a theoretical robustness bound is pursued, deriving it rigorously with explicit probabilistic assumptions rather than presenting an informal one.

## Data Availability

BraTS data are third-party and subject to their own data-use terms; they are not redistributed with this paper or its code release. Code, trained checkpoints, and all result files underlying this paper are available under the MIT license; see the citation record for version and access details. Code, trained checkpoints, and result files are available at https://github.com/ka-cyber/NeuroMesh, with the specific manuscript version identified by the corresponding Git commit.

## Notes

https://www.kaggle.com/datasets/awsaf49/brats2020-training-data

https://github.com/ka-cyber/NeuroMesh

### Competing Interest Statement

The authors have declared no competing interest.

## References

[1] N. Srivastava, G. Hinton, A. Krizhevsky, I. Sutskever, and R. Salakhutdinov, “Dropout: a simple way to prevent neural networks from overfitting,” J. Mach. Learn. Res., vol. 15, no. 1, pp. 1929–1958, 2014.

[2] Z.-H. Zhou, Ensemble Methods: Foundations and Algorithms. Boca Raton, FL, USA: CRC Press, 2012.

[3] D. Lepikhin, H. Lee-Thorp, C. Alberti, et al., “GShard: Scaling giant models with conditional computation and automatic sharding,” in Proc. Int. Conf. Learn. Represent. (ICLR), 2021.

[4] N. Shazeer, N. Parmar, J. Prabhu, et al., “Outrageously large neural networks: The sparsely-gated mixture-of-experts layer,” in Proc. Int. Conf. Learn. Represent. (ICLR), 2017.

[5] H. Liu, K. Simonyan, and Y. Yang, “DARTS: Differentiable architecture search,” in Proc. Int. Conf. Learn. Represent. (ICLR), 2019.

[6] Y. Gao, H. Yang, P. Zhang, C. Zhou, and Y. Hu, “Graph neural architecture search,” in Proc. 29th Int. Joint Conf. Artif. Intell. (IJCAI), 2020, pp. 1403–1409.

[7] J. Frankle and M. Carbin, “The lottery ticket hypothesis: Finding sparse, trainable neural networks,” in Proc. Int. Conf. Learn. Represent. (ICLR), 2019.

[8] A. Madry, A. Makelov, L. Schmidt, D. Tsipras, and A. Vladu, “Towards deep learning models resistant to adversarial attacks,” in Proc. Int. Conf. Learn. Represent. (ICLR), 2018.

[9] J. M. Cohen, E. Rosenfeld, and J. Z. Kolter, “Certified adversarial robustness via randomized smoothing,” in Proc. Int. Conf. Mach. Learn. (ICML), 2019.

[10] M. Lecuyer, V. Atlidakis, R. Geambasu, D. Hsu, and S. Jana, “Certified robustness to adversarial examples with differential privacy,” in Proc. IEEE Symp. Secur. Privacy (S&P), 2019, pp. 656–672.

[11] O. Ronneberger, P. Fischer, and T. Brox, “U-Net: Convolutional networks for biomedical image segmentation,” in Proc. Med. Image Comput. Comput.-Assisted Intervent. (MICCAI), 2015, pp. 234–241.

[12] T. N. Kipf and M. Welling, “Semi-supervised classification with graph convolutional networks,” in Proc. Int. Conf. Learn. Represent. (ICLR), 2017.

[13] Y. Bengio, N. Léonard, and A. Courville, “Estimating or propagating gradients through stochastic neurons for conditional computation,” arXiv:1308.3432, 2013.

[14] M. Havaei, N. Guizard, N. Chapados, and Y. Bengio, “HeMIS: Heteromodal image segmentation,” in Proc. Med. Image Comput. Comput.- Assisted Intervent. (MICCAI), 2016, pp. 469–477.

[15] I. Loshchilov and F. Hutter, “Decoupled weight decay regularization,” in Proc. Int. Conf. Learn. Represent. (ICLR), 2019.

[16] B. H. Menze, A. Jakab, S. Bauer, et al., “The multimodal brain tumor image segmentation benchmark (BRATS),” IEEE Trans. Med. Imag., vol. 34, no. 10, pp. 1993–2024, Oct. 2015.

[17] S. Bakas, H. Akbari, A. Sotiras, et al., “Advancing the cancer genome atlas glioma MRI collections with expert segmentation labels and radiomic features,” Sci. Data, vol. 4, Art. no. 170117, 2017.

[18] S. Bakas, M. Reyes, A. Jakab, et al., “Identifying the best machine learning algorithms for brain tumor segmentation, progression assessment, and overall survival prediction in the BRATS challenge,” arXiv:1811.02629, 2019.

